# Meta-transcriptomic analysis of companion animal infectomes reveals their diversity and potential roles in animal and human disease

**DOI:** 10.1101/2024.04.07.588491

**Authors:** Wei-Chen Wu, Yuan-Fei Pan, Wu-Di Zhou, Yu-Qi Liao, Min-Wu Peng, Geng-Yan Luo, Gen-Yang Xin, Ya-Ni Peng, Tongqing An, Bo Li, Huanle Luo, Vanessa R. Barrs, Julia A. Beatty, Edward C. Holmes, Wenjing Zhao, Yuelong Shu

## Abstract

Companion animals such as cats and dogs harbor diverse microbial communities that can potentially impact human health due to close and frequent contact. To better characterize their total infectomes and assess zoonotic risks, we performed meta-transcriptomic profiling on 239 samples from cats and dogs collected across China, comparing the similarities and differences between animal species (cats or dogs), sampling sites (rectal or oropharyngeal), and health status (healthy or diseased). We identified 24 viral species, 270 bacterial genera, and two fungal genera, including many known pathogens such as *canine parvovirus*, *Clostridium difficile*, and *Candida albicans,* as well as opportunistic pathogens such as *canine vesivirus*. Microbial compositions differed mainly according to sampling site (i.e., rectal and oropharyngeal swabs), and less so between host species and health status. Notably, we detected 27 potential zoonotic pathogens, such as *alphacoronavirus 1*, among all sampling sites, hosts, and health status, underscoring substantial zoonotic risks requiring surveillance. Overall, our meta-transcriptomic analysis reveals a landscape of actively transcribing microorganisms in major companion animals, including key pathogens, those with the potential for cross-species transmission, and possible zoonotic threats.

## Introduction

Cats and dogs are the most common companion animal species, with a global population size estimated to exceed 900 million^1,2^. Close contact between pet dogs and cats and their owners raises the possibility of microbial “sharing”, the impacts of which are unclear. On one hand, exposure to companion animals at an early age may provide immune benefits to human health^3-7^. However, living with companion animals might increase the risk of bidirectional transmission of some infectious diseases. Indeed, cats and dogs have been identified as potential sources of zoonoses of varying severity in humans, including viral pathogens (such as rabies virus and norovirus)^8-12^, bacteria (such as *Bordetella*, *Campylobacter* and *Salmonella*)^13-16^, fungi (such as dermatophytes)^17^, as well as parasites (such as *Toxoplasma gondii*)^18^, causing human illnesses ranging from mild rashes to severe nerve damage^8^. Additionally, companion animals may become infected with some human respiratory viruses, such as influenza viruses^19^, as well as multidrug-resistant bacteria, such as extended-spectrum beta-lactamase-resistant *Escherichia coli* and *Clostridium difficile*^12,20-26^, posing potential risks to human and animal health.

Recently, total RNA sequencing (i.e., meta-transcriptomics) has been used to simultaneously characterize the total “infectome” (i.e. all microbial pathogens within a sample, including bacteria, fungi and DNA/RNA viruses) of humans^27,28^, wildlife animals^29-31^, livestock species^32^, and arthropod vectors^33^. Not only does this method identify microbial taxa at the most precise taxonomic level possible, but it also enables accurate estimation of microbial abundance levels by measuring actively transcribing RNA molecules^28,32^. This also facilitates analysis of the interactions between pathogens and microbiota within a diseased system, as demonstrated by the characterization of microbial dysbiosis following SARS-CoV-2 infection in humans^27,34^.

Despite numerous metagenomic studies on cats and dogs, most previous studies have focused on characterizing the commensal bacterial microbiome^35-39^ or discovering novel pathogens^40-45^. In contrast, little is known about the entire infectome of companion animals. Herein, we used a meta-transcriptomics approach to perform in-depth comparisons of microbial diversity across different sampling sites (rectal/oropharyngeal swabs), species (dogs/cats), and health status (healthy/diseased), in doing so identifying microbial taxa associated with respiratory and gastrointestinal diseases. In addition, we further characterized the extent and pattern of cross-species microbial transmission among species, and assessed the risk that the microorganisms present in companion animals might pose to animal and human health.

### Results Study Design

We sampled a total of 1130 oropharyngeal and/or rectal swab samples from 571 companion animals – cats and dogs – across four provinces in China. Based on these samples, we set up a balanced experimental design for meta-transcriptomics analyses. This comprised 239 samples from diseased and healthy animals (122 : 117), cats and dogs (120 : 119), and oropharyngeal and rectal swabs (117 : 122) (Fig. 1). The diseased group included animals showing either respiratory (e.g., sneezing, coughing, nasal discharge) or gastrointestinal signs (e.g., diarrhea, vomiting) (61 : 61). Within the diseased group, rectal swabs were solely collected from animals with gastrointestinal signs, while oropharyngeal swabs were solely obtained from those with respiratory symptoms. Thus, comparisons between sampling sites in diseased animals in the following analyses implied a comparison between animals showing respiratory and gastrointestinal signs. For each combination (i.e., health status, animal species, and sampling sites), 27 to 34 individuals were used as replicates, the geographic distributions of which were balanced across the four Chinese provinces in which sampling took place (Fig. 1, Table S1).

**Figure 1.**
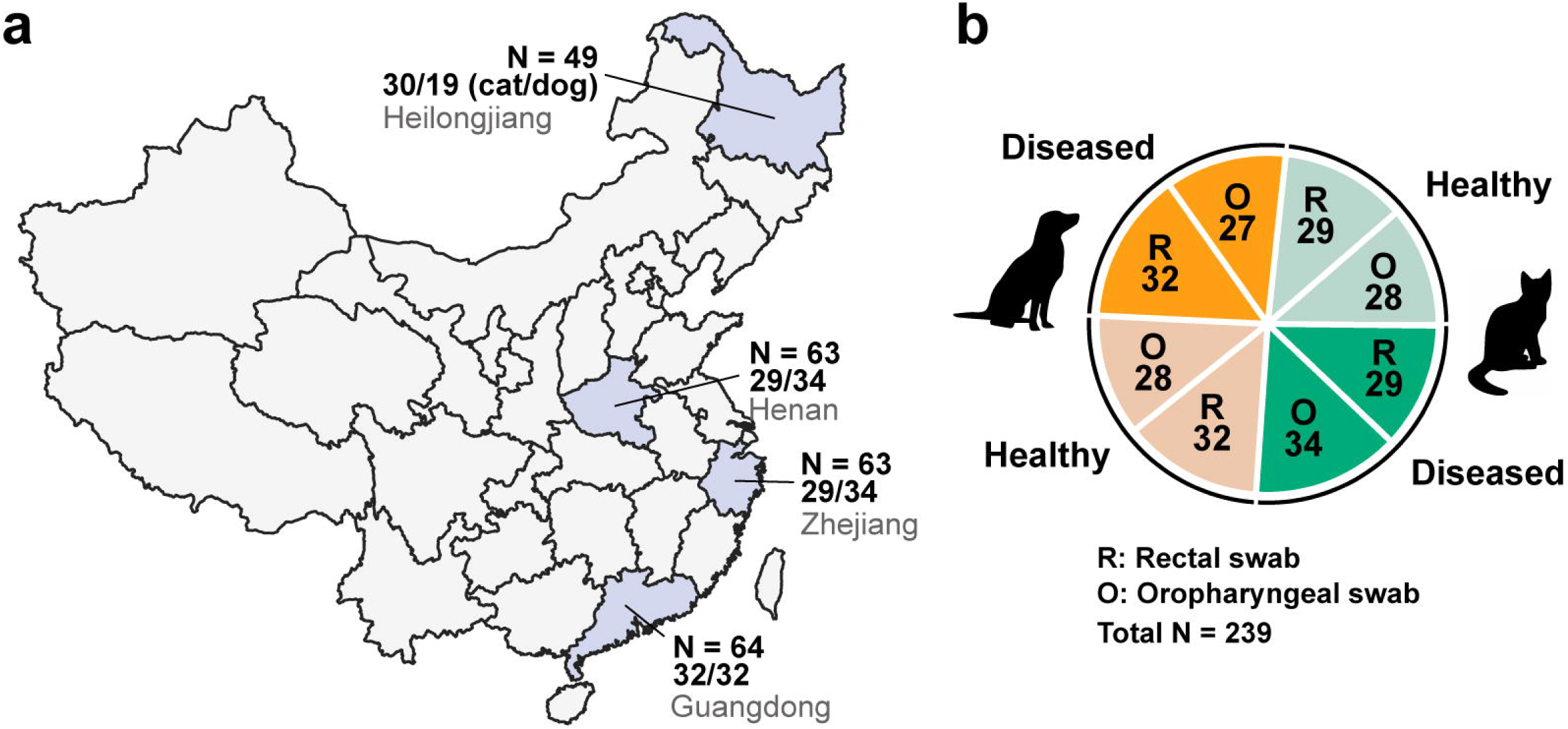
Sample overview. **(a)** Geographical distribution within China of the companion animal samples collected and the sample size at each location. **(b)** Sample types and corresponding sample sizes.

### Characterization of total infectome

Our meta-transcriptomic analysis of the total infectome within each sample revealed 24 known mammal-associated viral species, 270 bacterial genera, and two fungal genera (Fig. 2). The virus species identified here comprised 17 RNA viruses, six DNA viruses and one exogenous retrovirus species, in total representing 19 viral genera and 14 families (Fig. S1-2). Many of these are well-known pathogens in domestic animals, such as *alphacoronavirus 1*, *feline calicivirus*, *canine parvovirus*, *canine morbillivirus*, and *Norwalk virus*. We also found viruses of uncertain pathogenic potential that were present at high prevalence in our samples, including *canine vesivirus* (*N* = 27 animals), *canine kobuvirus* (*N* = 16), *canine astrovirus* (*N* = 14), and *mamastrovirus 2* (*N* = 13). Generally, an average of 1.61±1.04 (mean±s.d.) and up to five species of mammal-associated viruses were identified in each animal, with significantly more in diseased than healthy animals (1.87±1.19 in diseased animals, 1.24±0.58 in healthy animals, Wilcoxon test p = 0.0006; Fig. S3).

**Figure 2.**
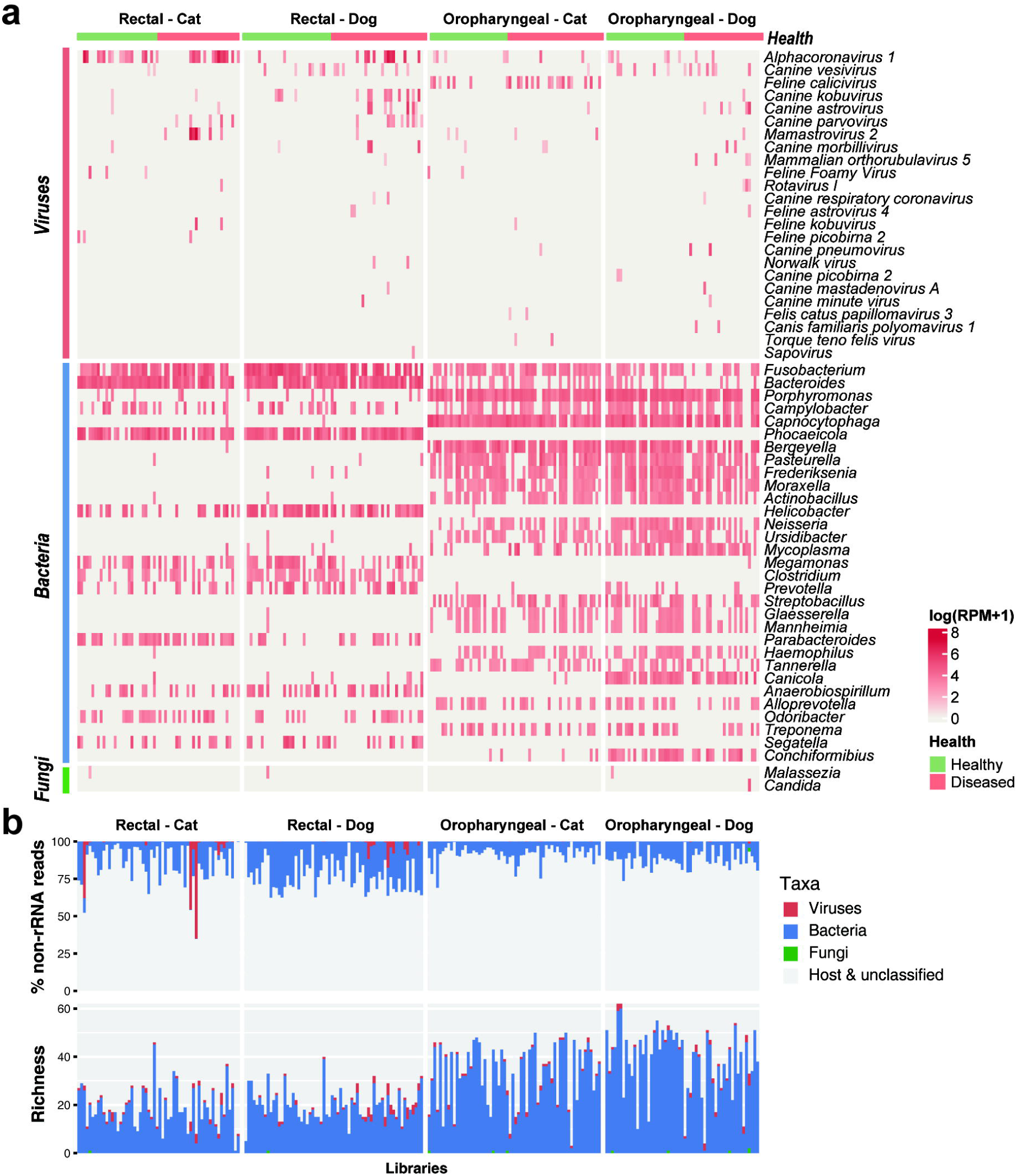
Overview of the infectomes of cats and dogs. **(a)** Heatmaps illustrating the microbial abundance in each individual sample. The abundance of viruses is quantified at the species level, while bacterial and fungal abundance are quantified at a genus level resolution. For clarity, bacterial genera with <40 positive samples were omitted from the visual representation. **(b)** The abundance and diversity of viruses, bacteria and fungi in each library. The diversity of viruses was quantified by species richness (i.e., the number of viral species per sample), while the diversity of bacteria and fungi was quantified by the number of genera.

The bacterial genera identified here largely comprised the normal flora of cats and dogs, including those at very high prevalence and abundance (i.e., up to 67% of total non-rRNA reads), such as *Fusobacterium*, *Bacteroides*, *Phocaeicola* and *Helicobacter* in the rectal swab samples, and *Porphyromonas*, *Capnocytophaga* and *Bergeyella* in the oropharyngeal swabs (Fig. S4). The dominant bacterial genera differed between rectal and oropharyngeal swabs, but were similar between cats and dogs (Fig. S4). Of note, we identified 23 bacterial species that can infect humans, including *Clostridioides difficile*, *Campylobacter jejuni*, *Moraxella catarrhalis, Capnocytophaga cynodegmi*, and *Helicobacter canis.* (Fig. S5; Table S2).

Two fungal species were identified in this study – *Candida albicans* was identified in the oropharyngeal swab sample of a single dog with respiratory symptoms, while *Malassezia globosa* was detected in three healthy animals (two dogs and one cat). Importantly, both species are associated with opportunistic infections of the human skin, respiratory tract, and genitourinary tract, suggesting zoonotic potential.

Overall, based on meta-transcriptomic sequencing, the total infectome accounted for an average of 11.8% (0.02%-65.22%) of total non-rRNA reads (Fig. 2b), many of which belonged to commensal bacteria (11.3%; range 0.01%-37.66% of total reads). In comparison, the mammal-associated viruses were much less abundant (median 0.016%, range 4.8×10^-5^%-65.22%), although in five of the samples the number of viral reads exceeds 10% of the total reads.

### Comparisons of microbial composition and diversity

We compared overall microbial diversity across different sampling site (oropharyngeal/rectal swabs), animal species (dogs/cats), and health status (healthy/diseased) (Fig. 3A, Fig. S3). Oropharyngeal swabs had significantly greater bacterial richness than rectal swabs (Wilcoxon test, p<0.001). Oropharyngeal samples from dogs exhibited higher bacterial richness than those of cats (p<0.001), while cat rectal samples showed higher viral richness compared to dogs (p=0.005). With respect to health status, bacterial richness was higher in oropharyngeal samples from healthy compared to sick dogs (p=0.001). Conversely, viral richness was significantly higher in oropharyngeal swabs of diseased animals (cat oropharyngeal swabs, p=0.034; dog oropharyngeal swabs, p=0.013) and in rectal swabs of dogs (p<0.001). Average viral abundance levels were also significantly higher in diseased animals (p<0.05). Bacterial and viral community composition (beta diversity) differed significantly among sampling sites for both bacteria (PERMANOVA test, R^2^=0.26, p<0.001) and viruses (R^2^=0.038, p<0.001). For bacteria, the composition differed between cats and dogs in both swabs (p<0.001), and between health status in oropharyngeal swabs (p=0.005). Viral composition was significantly different between species (rectal swabs, p=0.003; oropharyngeal swabs, p < 0.001) and health status (rectal swabs, p=0.002; oropharyngeal swabs, p = 0.009) in both swabs.

**Figure 3.**
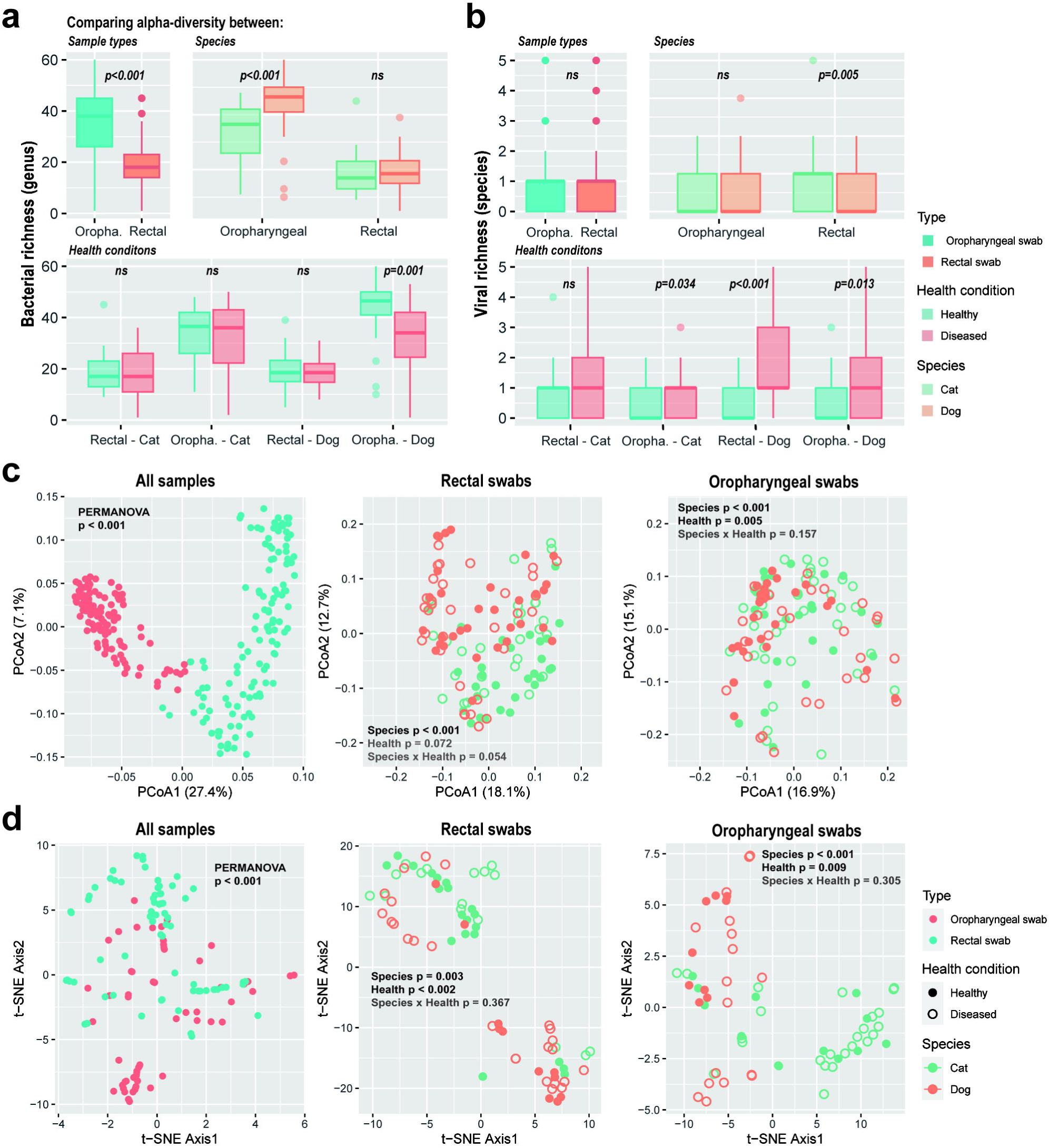
Comparisons of alpha and beta diversity among species, sample type and health conditions. **(a, b)** Comparison of bacterial richness (a; number of bacterial genera per sample) and viral richness (b; number of viral species per sample) among sample types (anal/throat swabs), species (cats/dogs) and health conditions (healthy/diseased). **(c, d)** Comparison of bacterial genera compositions (c) and viral species compositions (d) among samples. P-values from PERMANOVA tests are shown at the top. For all samples, we only tested the effect of sample type. For specific sample types we tested the effect of species, health condition and their interaction. Significant results are shown in bold black fonts.

### Possible disease associations

Viral species potentially associated with respiratory and/or gastrointestinal diseases were inferred by comparing their prevalence among healthy and diseased animals showing either respiratory or gastrointestinal signs (Fig. 4). Of note, within the diseased group rectal swabs were solely collected from animals with gastrointestinal signs, while oropharyngeal swabs were solely obtained from those with respiratory symptoms. Thus, comparisons between sampling sites in diseased animals also imply a comparison between animals showing respiratory and gastrointestinal signs. Many viruses were only associated with diseased animals, such as *canine parvovirus, canine astroviruses,* and *mammalian orthorubulavirus 5*, and also showed specificity to particular sampling site or animal species. All other viruses were detected in both healthy and diseased animals, although some were at greater prevalence levels in diseased than healthy animals, such as *feline calicivirus* and *canine morbillivirus* in dog samples. Interestingly, in the case of *alphacoronavirus 1*, disease associations might differ between sampling site and animals. Indeed, this virus was only associated with disease in dog rectal samples (Fig. 4).

**Figure 4.**
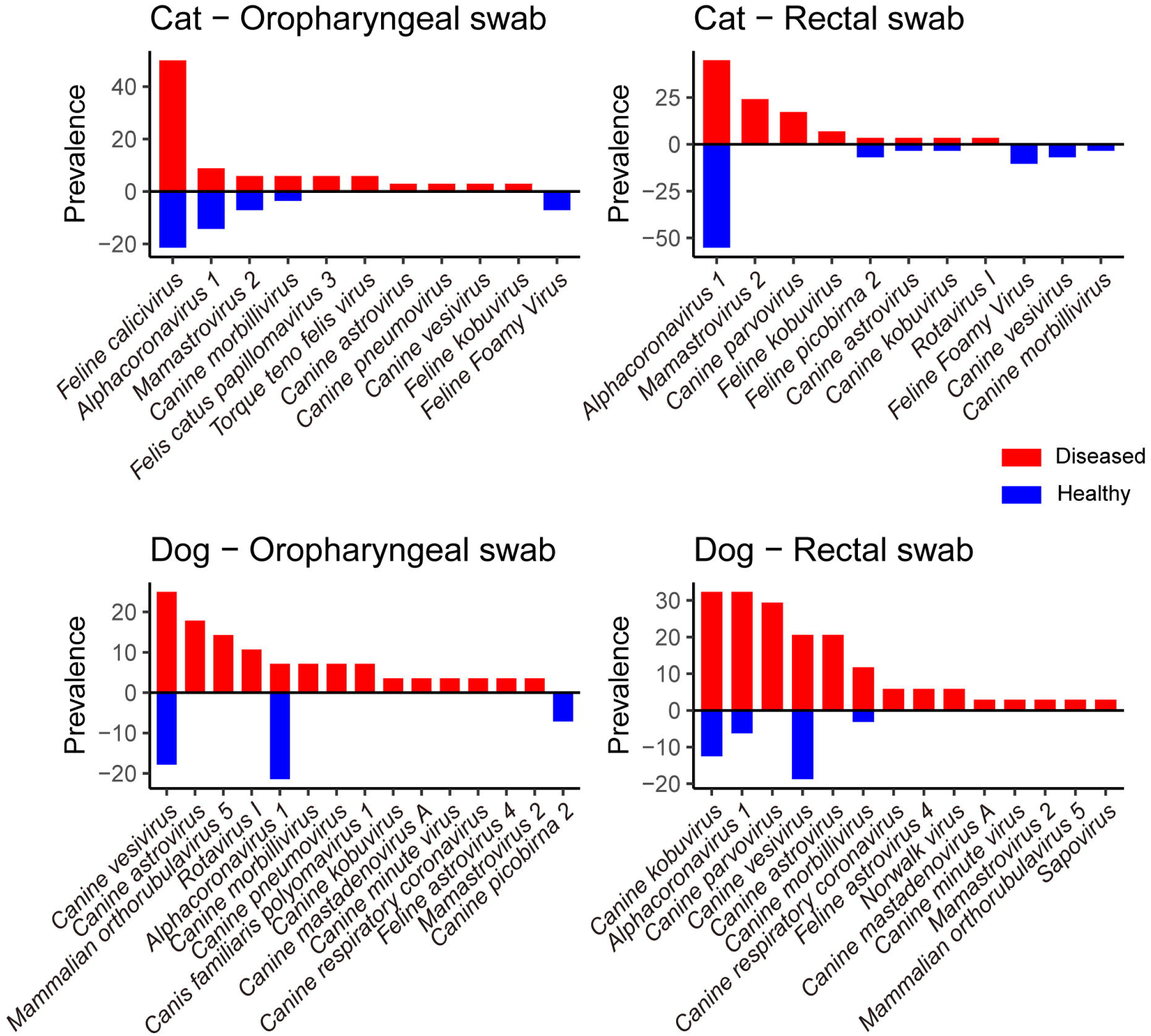
Differences in prevalence in viral species among healthy and diseased animals. The prevalence of viral species detected in specific samples, sorted by prevalence in diseased animals (red) and then by prevalence in healthy animals (blue).

Because potential pathogens may show significantly greater abundance in diseased animals, we performed differential abundance analysis between healthy and diseased animals using three independent tests – DESeq2, LefSe, and Wilcoxon tests. Only microbial species/genera with more than five positive samples were included in this analysis. This analysis showed that six virus species and 26 bacterial genera were associated with disease (Fig. 5, Fig. S6). In contrast, many bacterial genera showed the opposite trend: higher abundance in healthy than diseased (Fig. 5, Fig. S6).

**Figure 5.**
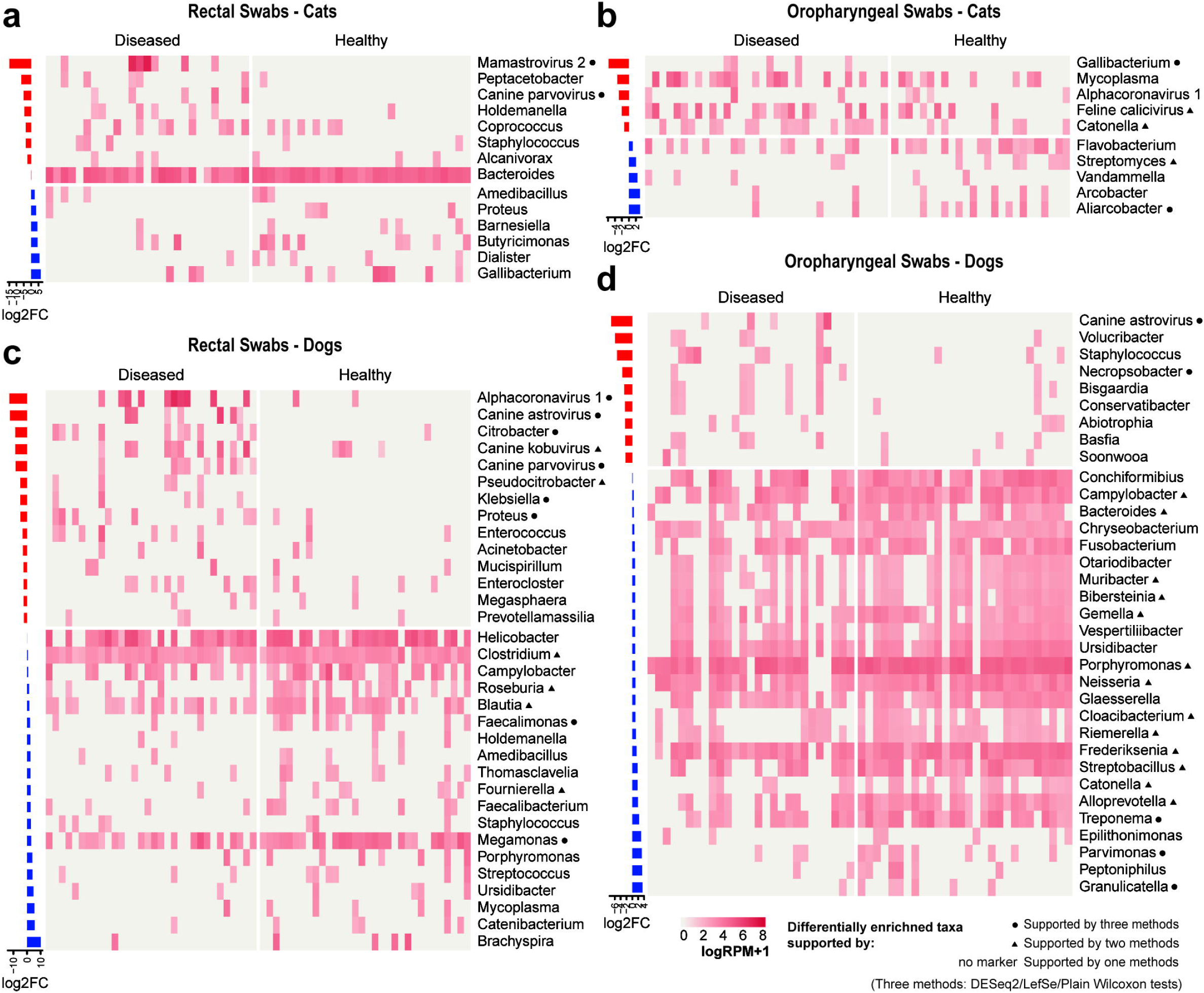
Differential abundance analysis of the infectomes of healthy and diseased animals. **(a-d)** Heatmaps demonstrating the abundance differences of microbial taxa (bacterial and fungal genera; viral species) among diseased and healthy animals. Differential abundance analyses (DAA) were performed using three methods, LEfSe, DESeq2 and Wilcoxon tests. The bar plots on the left exhibit the log-transformed fold change in abundance (RPM) of the corresponding taxa.

Differences in pathogenic potential and tissue/host preference were also apparent at the genotype/lineage level as shown in intra-specific phylogenetic trees of several potentially pathogenic and a non-pathogenic virus species (Fig. 6). For *alphacoronavirus 1*, sequences from the canines-associated lineages were primarily found in sick dogs, whereas those from the feline-associated lineage were found in both healthy and diseased cats. Similarly, distinct lineages were associated with differences in pathogenicity for feline calicivirus, with certain lineages predominantly associated with diseased cats, while another lineage was associated with both healthy and diseased cats, suggesting opportunistic infections. In contrast, no such patterns were observed for *canine vesivirus* (Fig. 6).

**Figure 6.**
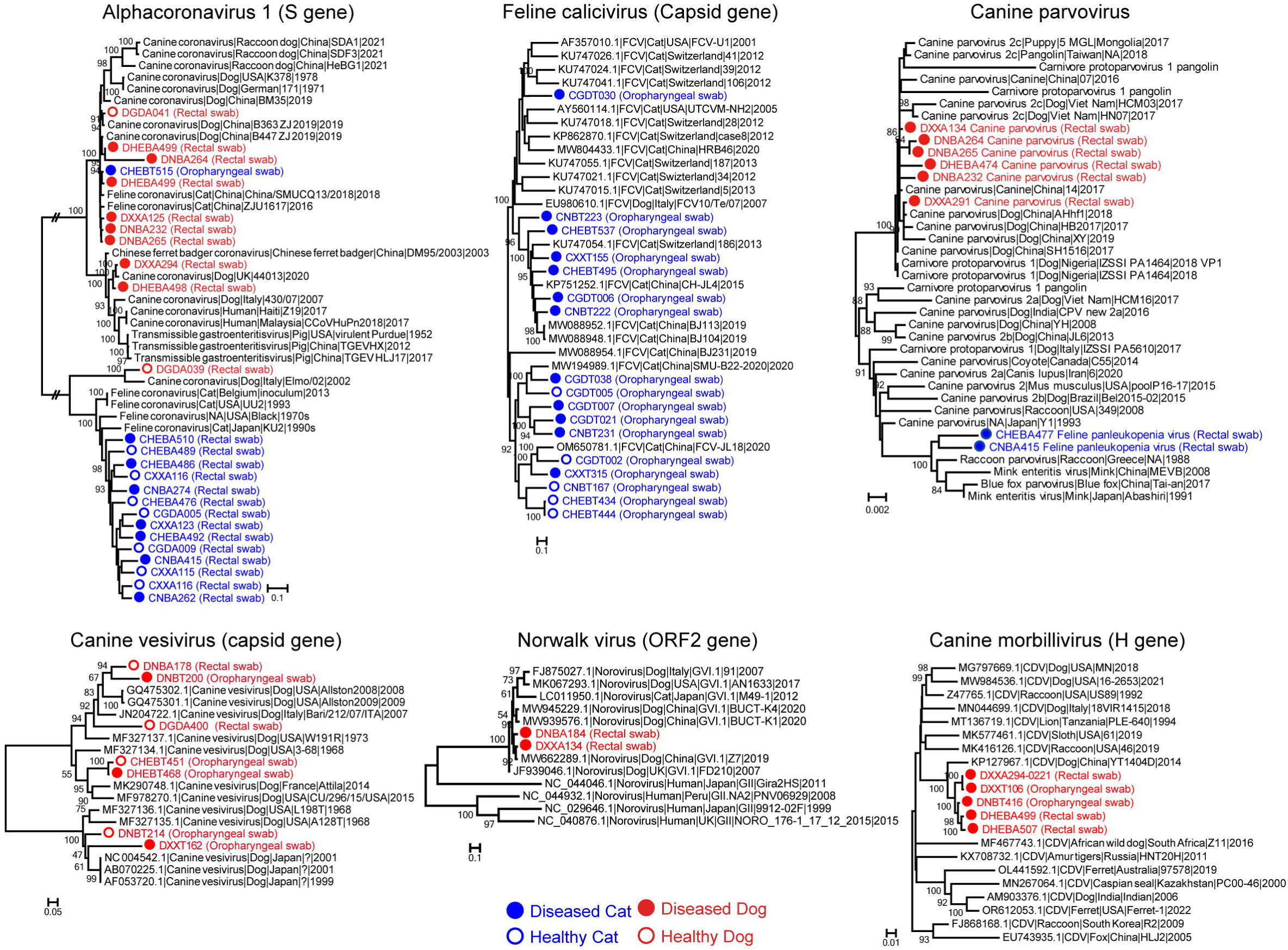
Maximum likelihood phylogenetic trees of six viral pathogens. The phylogenies were inferred using key functional genes (nucleotide sequences): Alphacoronavirus 1 (S gene), Feline calicivirus (capsid gene), Carnivore protoparvovirus 1 (VP1 gene), Nowalkvirus (ORF2 gene), Canine vesivirus (capsid gene) and Canine distemper virus (H gene). Host species and tissue type are indicated. The trees are midpoint rooted for clarity, with branch lengths reflecting the number of substitutions per site.

### Shared infectious agents and zoonotic risks

We identified six viral species that infected both cats and dogs, suggesting a history of cross-species transmission (Fig. 7). All six virus species were detected in rectal swabs of dogs and cats, with the exception that *canine morbillivirus* was not found in cat rectal swab. Four viral species (*canine morbillivirus and alphacoronavirus 1, canine vesivirus and canine astrovirus*) were additionally detected in oropharyngeal swabs. Of note, *alphacoronavirus 1* appeared at high prevalence in both species and sampling sites, suggesting a broad host range and tissue tropism (Fig. 6). Additionally, these data revealed a number of mixed infections (Fig. S7, Table S3). For instance, *alphacoronavirus 1* frequently co-occurred with *canine parvovirus* and *canine kobuvirus* (Chi-squared tests, p < 0.05), and significant positive correlations among their abundance were also detected (Spearman tests, p < 0.05).

**Figure 7.**
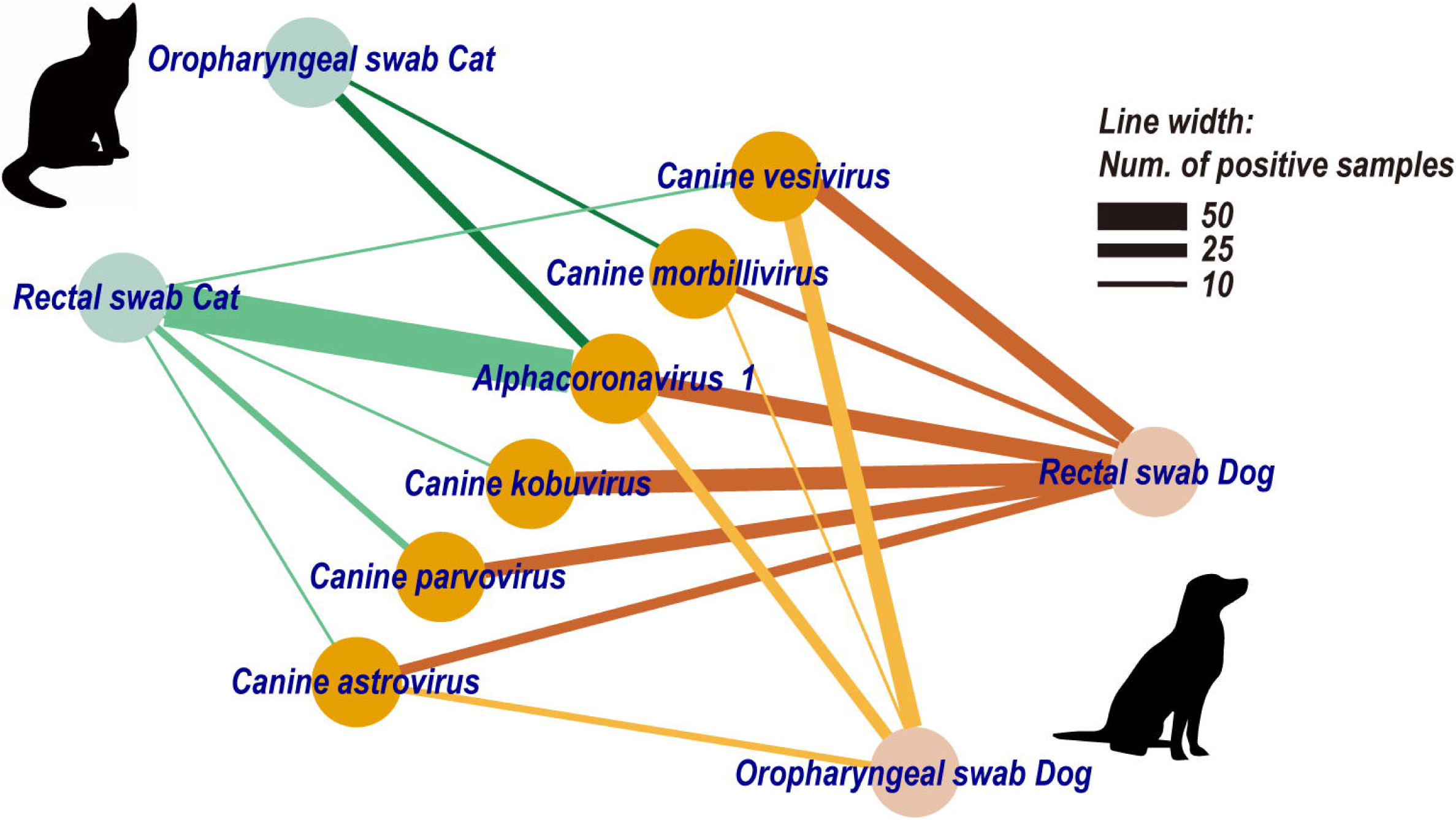
The virus sharing network between cats and dogs. Nodes represent either viral species or specific sample types (rectal/throat swabs of cats/dogs). Lines between nodes indicate the presence of a specific viral species in corresponding samples, and the line width is proportional to the number of positive samples.

Notably, we identified 27 known (opportunistic) zoonotic pathogens to humans, comprising two viral species, 23 bacterial species, and two fungal species (refer to Table S2). Of these, seven species were significantly more prevalent in rectal swabs (Chi-square test, p < 0.05), whereas another eight species were more prevalent in oropharyngeal swabs. Many of these zoonotic pathogens (63%, 17 species) were detectable in both dogs and cats, with no statistically significant differences in their prevalence between the two host species (Chi-square tests, p < 0.05). A similar pattern was observed across health status, where 62% of these pathogens (17 species) were detectable in both healthy and diseased animals.

## Discussion

The infectomes of companion animals such as cats and dogs may have important implications for animal and perhaps human health^46^. While previous studies have characterized infectomes through the metagenomic sequencing of total DNA (e.g., ref.^36,47^), our study employed a meta-transcriptomic approach to specifically profile the actively transcribing portion of the infectome. This enabled comprehensive characterization of viruses as well as bacteria and fungi.

Our analysis revealed that bacteria dominated the healthy pet infectome, comprising an average 11.3% of total non-rRNA reads. The predominant bacterial genera identified, such as *Fusobacterium, Bacteroides, Phocaeicola*and *Helicobacter* in rectal swabs, and *Porphyromonas, Capnocytophaga* and *Bergeyella* in respiratory swabs, were largely consistent with previous studies^48-52^. This implies that overall transcriptional activity correlates with genome abundance for dominant bacteria. Also in accord with previous studies, our results showed that microbiome of healthy animals differs between tissue types^48,53^, but less so between different host species^48^. Some specific bacteria species/genera can be detected in both cats and dogs, and there is evidence suggesting that some of these may be zoonotic, causing diseases from diarrhea (*Clostridioides difficile^22^, Fusobacterium varium^54^),* to serious bite-wound infections (*Pasteurella spp.^55,56^, Capnocytophaga spp.^57^, and Neissieria spp*.^58^).

Although fungi are considered part of the normal mycobiota in animals^59^, metagenomic studies suggest that their abundance is much lower than bacteria, and they were rarely detected in this study. For instance, in a study on dog intestines, fungi comprised only 1% of the microbial DNA reads, while bacteria accounted for 98%^60^. This is consistent with the notion that bacteria dominate the microbiome. Given the low abundance of most fungi, the actively transcribing portion might be even lower. While meta-transcriptomics is still capable of detecting abundant, actively transcribing fungi, which could indicate ongoing infections (e.g., candidiasis), it is important to note that the RNA extraction methods employed in this study did not involve fungal cell disruption techniques such as bead-beating, which are essential for enriching fungal DNA in metagenomics studies^61^.

For the viral component of infectome, we focused specifically on mammal-associated viruses with likely direct relevance to animal and human health. RNA viruses dominated this component in terms of both diversity and abundance. In contrast to bacteriome which is dominated by commensal species, the pet virome consists of many well-known pathogens such as *feline calicivirus* and *canine parvovirus*, as well as emerging viral pathogens like *canine vesivirus^62^*, *canine kobuvirus^63^*. While viruses exhibited lower overall abundance than bacteria, viral reads exceeded 10% of the total in some samples, indicating that the virome represents an important part of the active infectome. Notably, viral composition differed less between sampling sites (rectal or oropharyngeal) than bacteria, likely the result of systemic infection (which is the case for *canine morbillivirus^64^*, etc.). Furthermore, our results documented the sharing of several viral species between cats and dogs, including viral species that pose potential zoonotic risks to humans such as *alphacoronavirus 1*, highlighting the public health importance of monitoring pets using meta-transcriptomics.

Comparative analysis between healthy and diseased animals provided further evidence on the potential pathogenic roles of several viruses with previously ambiguous disease associations, such as *canine astrovirus^40^*. In particular, their enrichment in diseased compared to healthy animals suggests that they may play a role as opportunistic pathogens. Other viruses, including *canine vesivirus*, showed high prevalence regardless of health status, suggesting that they are normal components of the virome that may become pathogenic in contexts of disrupted homeostasis. While differential abundance testing identified taxa enriched in disease, limited sample sizes and genus-level resolution (bacteria and fungi) could have obscured detection of some pathogens.

Importantly, we detected a number of zoonotic viral, bacterial and fungal pathogens that might cause opportunistic infections in humans, including *alphacoronavirus 1^65^, Norwalk virus^66^*, *Clostridioides difficile^22^*, *Campylobacter jejuni^11^* and *Candida albicans^67^.* Their presence in these companion animals highlights potential risks of disease transmission to humans, especially immunocompromised individuals or those with frequent animal contact. Comprehensive surveillance of pets and their owners is therefore warranted to better understand zoonotic risks. Importantly, our results revealed that while these potential zoonotic pathogens exhibited distinct sampling site preference, 22% (6 species) were detectable in both sampling sites. Strikingly, 62% of these zoonotic pathogens were detectable in both healthy/diseased dogs and cats. These findings underscore the importance of surveillance in both dogs and cats, regardless of their health status, while targeting specific sampling sites for detecting intended pathogens.

Interestingly, our analysis identified fewer zoonotic viral species (two species) than zoonotic bacterial species (23 species) in companion animals. This contrasts with studies of other domestic animals such as pigs, where numerous zoonotic viruses have been characterized. For example, metagenomic analyses of pigs^32^ have revealed the presence of zoonotic viruses like influenza A virus, hepatitis E virus, and multiple arbovirus species (e.g., Japanese encephalitis virus, Getah virus and Zika virus) in addition to common zoonotic bacterial pathogens such as *Streptococcus suis*. The relatively lower viral zoonotic risk observed here for companion animals may be attributable to differences in viral host range, tropism and transmission dynamics between companion animals versus livestock species. Nonetheless, the zoonotic viral threats identified again highlights the importance of viral surveillance even in companion animal populations.

In summary, our meta-transcriptomic study reveals the landscape of actively transcribing microbes that constitute the infectomes of companion animals. We identified key pathogenic viruses, bacteria and fungi, elucidated their distribution among host species, tissue types and health status, and evaluated their zoonotic potential.

## Methods

### Ethics

This research, including the procedures and protocols of specimen collection and processing, was reviewed and approved by the Institutional Animal Care and Use Committee, Sun Yat-sen University (No. SYSU-IACUC-MED-2021-B0123).

### Sample Collection

Between March 2021 to March 2022, we sampled domestic animals (i.e., cats and dogs) from pet hospitals and pet care centers located in five major cities distributed throughout China: Guangzhou (Southern China), Shenzhen (Southern), Xinxiang (Central), Ningbo (Eastern) and Harbin (Northern). Both healthy animals and diseased animals were enrolled. The diseased animals had either respiratory (e.g., sneezing, coughing, nasal discharge) or gastrointestinal symptoms (e.g., diarrhea, vomiting, loss of appetite) after the initial diagnosis by the veterinarians who carried out the sampling. Healthy animals showed no clinical symptoms at the time of sampling and were taken to the facility for health check, cleaning, and beauty services. For each animal, oropharyngeal or rectal swab sampling was performed depending on the clinical symptoms. The samples were immediately immersed in virus preservation solution (SHENQI Biotech), placed on dry ice, and then transferred to -80°C refrigerator for storage. Detailed sample information is provided in Supplementary Table S1.

### Sample processing and meta-transcriptomics analyses

Total RNA was extracted from 239 individual swab sample using the RNeasy Plus universal mini kit (QIAGEN) according to the manufacturer’s instructions. The concentration of extracted RNA was determined using a Qubit™ RNA High Sensitivity Kit (Thermo Fisher Scientific). RNA sequencing library construction were performed using the Trio RNA-Seq Library Preparation Kit (NuGEN Technologies) which targeted low-concentration RNA as starting material and used an AnyDeplete probe (NuGEN Technologies) to remove host ribosomal RNA. The concentration and quality of constructed library were determined using Qubit™ dsDNA Quantification Assay Kit (Thermo Fisher Scientific) and Qsep100 (Bioptic), respectively. Paired-end (150 bp) sequencing of the libraries was performed on the Illumina NovaSeq platform. The quality control of subsequent sequencing reads was performed using BBDuk (version 38.62) (downloaded from https://jgi.doe.gov/data-and-tools/software-tools/bbtools/). rRNA reads were removed by mapping against a comprehensive rRNA reference sequence collection downloaded from the SILVA database^68^ (https://www.arb-silva.de/) using bowtie2 (version 2.3.5.1)^69^. The deduplication of the reads was performed using CD-HIT (version 4.8.1)^70^, and the processed reads were then *de novo* assembled into contigs using MEGAHIT (version 1.2.8)^71^ for subsequent microbial identification and profiling.

### Profiling vertebrate viruses

We identified potential RNA and DNA viruses by comparing the assembled contigs against a NCBI reference virus database using blastn^72^ (version 2.9.0) and a NCBI non-redundant protein database (nr) using Diamond blastx^73^ (version 0.9.25). The resulting viral contigs were subsequently compared to the non-redundant nucleotide database (nt) to identify and remove contigs or regions in the contigs that were related to host or bacterial genomes. The remaining viral contigs were categorized to the species level based on sequence homology with reference viral genomes as well as that with each other. Novel viral species were defined based on the guidelines provided by International Committee on Taxonomy of Viruses (ICTV).

Among all the viral species identified, we only characterized the likely mammal-associated virome, with the component viruses determined based on whether they fell within well-defined vertebrate-specific viral genera or families following phylogenetic analysis^74^. The quantification of each viral species was carried out by mapping non-rRNA reads to the viral genomes identified in this study as well as to related reference sequences. Read counts mapped to each sequence were summed at the species level. Index-hopping filtering was performed on the read count matrix using a threshold of 0.1%. The filtered read count matrix was then normalised by total non-rRNA read count to generate a RPM table.

### Profiling the composition of bacteria and fungi

We identified bacterial and fungal genera and quantified their abundance by mapping non-rRNA reads to a set of reference genomes. The reference genomes of bacteria were retrieved from the Genome Taxonomy Database^75^ (GTDB) release 214.1, which is a widely used database of curated bacteria genomes and taxonomy. For fungi, we used all representative fungal genomes from the NCBI Genome Assembly database. Before mapping reads to genomes, rRNA regions in these genomes were masked with the ambiguous nucleotide code “N” to reduce false-positive alignments to the highly-conserved rRNA gene. This was done by performing a blastn^72^ search against SILVA rRNA database^68^ (release 138) to locate rRNA and then mask them with a custom python script. When the reference genomes were prepared, non-rRNA reads were aligned against these genomes using bowtie2^69^. The sensitivity of the read alignment process was set to “—very-fast”. To reduce false-positives, we scanned the consensus sequences from read alignments for a set of universal marker genes using BUSCO v5.5.0^76^. If none of the marker genes were present in a consensus sequence, the whole alignment was considered as false-positive and removed. After these steps, read counts mapped to each genome were summed by genus, and index-hopping filtering was performed on the genera-level read count matrix as described above. The filtered read count matrix was then normalized by total non-rRNA read count to generate a RPM table. Bacterial and fungal genera at low abundance (RPM < 100) were removed.

### Identification of potential zoonotic pathogens

To identify potential zoonotic pathogens, we conducted species-level profiling for several bacterial and fungal genera known to harbor human pathogens (viruses were already identified to the species level; see above). A list of these genera was obtained from the KEGG DISEASE database^77^. We extracted two phylogenetically informative genes – rpoB^78^ and gyrB^60^ – from the consensus sequences of these genera generated during read mapping (as described above). The nucleotide sequences of the rpoB and gyrB genes were aligned with reference sequences of specific genera using MAFFT^79^. Phylogenetic trees were then estimated using the maximum likelihood method in IQ-TREE^80^ (employing the GTR+I+F+Γ_4_ substitution model). Clustering of the taxa generated here to known species was visually inspected on the phylogenies. Additionally, we calculated the average nucleotide identity (ANI) between the mapped consensus sequence and reference genomes. The identification of zoonotic pathogens to species resolution was supported by both ANI > 97% and evidence from rpoB or gyrB phylogeny.

### Genetic diversity of the pet infectome

We assessed the alpha diversity of the pet infectome by examining the richness of bacterial genera and viral species richness per sample. Due to low fungal diversity in our samples, these were omitted from the analyses of both alpha and beta diversity. We employed Wilcoxon rank-sum tests to compare the mean richness per sample across different sampling sites (oropharyngeal swabs vs. rectal swabs), species (cats vs. dogs), and health status (healthy vs. diseased). Statistical significance was determined at a threshold of p < 0.05. Notably, in the comparisons between sampling sites and species, only samples from healthy animals (n=117) were included.

Subsequently, we visualized the beta diversity of the pet infectome using ordination plots depicting microbial community compositions. For bacteria, we utilized Bray-Curtis distance metrics for beta diversity characterization and principal coordinate analysis (PCoA) for visualization. For viruses, Bray-Curtis distance was similarly utilized, but visualization was performed using t-distributed stochastic neighbor embedding^81^ (t-SNE), as the viral abundance matrix (viral species-by-sample) exhibited significant sparsity. To assess differences in pet microbial composition across sampling sites, species, and health status, we conducted permutational multivariate analysis of variance (PERMANOVA) using the adnois2 function^82^ within the R package vegan. Statistical significance was determined based on 999 permutations to generate p values.

### Differential Abundance Analyses (DAA)

To identity microbial taxa potentially associated with diseases in dogs and cats, we utilized differential abundance analyses (DAA). We performed three independent analyses using different methods of DAA – DESeq2^83^, LefSe^84^ and Wilcoxon tests – to produce as robust results as possible. These analyses were performed separately for each type of sample (i.e., dog rectal swab, dog oropharyngeal swab, cat rectal swab, and cat oropharyngeal swab). Microbial taxa with less than five positive samples were excluded from the analyses.

For the DESeq2 analysis, we used the abundance matrix (RPM values, microbial taxa-by-sample) as input. Significant microbial taxa were identified based on an adjusted p-value of < 0.05 and a log2FC (log-transformed fold change in read count between groups) exceeding 1 or below -1. For LefSe analysis, the abundance matrix (RPM values) was used as input. Results with LDA scores > 2 and p values < 0.01 were deemed significant (as recommended by the software). We performed Wilcoxon tests on each microbial taxon to compare their abundance (log-transformed RPM values) between disease and healthy pets. Any result with a p-value < 0.05 was deemed significant. Differential abundance analyses may produce different results, and those taxa supported by more than one method are thought to be more robust^85^. Finally, the differentially enriched taxa identified by the above methods were visualized by a heatmap.

### Phylogenetic analyses

We estimated the evolutionary history of six (potential) viral pathogens: *alphacoronavirus 1, feline calicivirus, canine parvovirus, Norwalk virus, canine vesivirus* and *canine morbillivirus*. Accordingly, phylogenetic trees were inferred based on key functional genes of each virus: the S gene for *alphacoronavirus 1*, capsid gene for *feline calicivirus*, VP1 gene for *canine parvovirus,* ORF2 gene for *Norwalk virus,* capsid gene for *canine vesivirus* and H gene for *canine morbillivirus*. We used IQ-TREE^80^ (version 2.2.0.3) to estimate phylogenies assuming a GTR+I+Γ_4_ model of nucleotide substitution. Bootstrap resampling (1000 replications) using ultra-fast bootstrapping option (-bb 1000) was used to assess the robustness of individual nodes.

We similarly performed phylogenetic analysis on bacteria. To retrieve marker gene sequence of bacteria, we aligned reads to representative genomes of several candidate genera (genera which may contain zoonotic or pathogenic species) using bowtie2 (version 2.5.0)^69^ and extracted rpoB gene (RNA polymerase beta subunit) from the consensus sequences. We then aligned these extracted nucleotide sequences together with reference sequences from GTDB release 214.1^75^ using MAFFT v7.515 ^79^ (L-INS-i algorithm, max iteration set to 1000). Phylogenetic trees were estimated using IQ-TREE (version 2.2.0.3)^80^, again employing the GTR+I+Γ_4_ substitution model.

## Supporting information

Suppmental Figure S1-S7

## Data availability

The meta-transcriptomic sequencing reads generated in this study have been deposited in the China National GeneBank DataBase (CNGBdb) under project accession code CNP0005429 (https://db.cngb.org/search/project/CNP0005429/).

## Code availability

Codes are available at: https://github.com/Augustpan/pet-infectome.

## Acknowledgements

This study was funded by grants from the National Key R&D Program of China (2022YFC2303801), Fund of Shenzhen Key Laboratory (ZDSYS20220606100803007), and Shenzhen Science and Technology Program (KQTD20200820145822023 and KCXFZ20211020172545006). E.C. Holmes was supported by an NHMRC (Australia) Investigator Award (GNT2017197) and by AIR@InnoHK administered by the Innovation and Technology Commission, Hong Kong Special Administrative Region, China.

We gratefully thank Yan-Xia Zhang, Yu-Tao Wu, Ying-Hong Guan, Zu-Qing Shi for their assistance in specimen collection. And we acknowledge all clinicians involved in the diagnosis of diseased animals and specimen collection.

## Author Contributions

Conceptualization, W-CW and VRB, JAB, ECH, WJZ and YLS; Methodology, W-CW, Y-FP, W-DZ, and ECH; Sample Collection and Processing, W-CW, W-DZ, Y-QL, M-WP, G-YL, G-YX, Y-NP, TQA and HLL; Data analysis, W-CW, Y-FP, W-DZ, Y-QL, and M-WP; Writing – Original Draft, W-CW and Y-FP; Writing – Review and Editing, W-CW, Y-FP, W-DZ, Y-QL, M-WP, G-YL, G-YX, Y-NP, TQA, BL, HLL, VRB, JAB, ECH, WJZ, and YLS. Funding Acquisition, YLS; Resources (sampling), W-CW, TQA, YLS; Resources (computational), W-CW and Y-FP; Supervision, BL, VRB, JAB, ECH, WJZ and YLS.

## Notes

### Competing Interest Statement

The authors have declared no competing interest.

## References

1 Li, L. et al. Viruses in diarrhoeic dogs include novel kobuviruses and sapoviruses. Journal of General Virology 92, 2534–2541 (2011).

2 Li, Y. et al. Virome of a feline outbreak of diarrhea and vomiting includes bocaviruses and a novel chapparvovirus. Viruses 12, 506 (2020).

3 Nermes, M. et al. Perinatal pet exposure, faecal microbiota, and wheezy bronchitis: is there a connection? International Scholarly Research Notices 2013 (2013).

4 Lødrup Carlsen, K. C., et al. Does pet ownership in infancy lead to asthma or allergy at school age? Pooled analysis of individual participant data from 11 European birth cohorts. (2012).

5 Havstad, S. et al. Effect of prenatal indoor pet exposure on the trajectory of total IgE levels in early childhood. Journal of Allergy and Clinical Immunology 128, 880–885. e884 (2011).

6 Azad, M. B. et al. Infant gut microbiota and the hygiene hypothesis of allergic disease: impact of household pets and siblings on microbiota composition and diversity. *Allergy*, Asthma & Clinical Immunology 9, 1–9 (2013).

7 Nermes, M., Endo, A., Aarnio, J., Salminen, S. & Isolauri, E. Furry pets modulate gut microbiota composition in infants at risk for allergic disease. Journal of Allergy and Clinical Immunology 136, 1688–1690. e1681 (2015).

8 Zhang, N., Song, C., Tao, X. & Zhu, W. Epidemiologic Features of Human Rabies in China from 2015-2021. Zoonoses 3 (2023).

9 Gelgie, A. E., Cavalerie, L., Kaba, M., Asrat, D. & Mor, S. M. Rabies research in Ethiopia: A systematic review. One Health 15, 100450 (2022).

10 Stull, J. W., Brophy, J. & Weese, J. Reducing the risk of pet-associated zoonotic infections. Cmaj 187, 736–743 (2015).

11 Gras, L. M. et al. Increased risk for Campylobacter jejuni and C. coli infection of pet origin in dog owners and evidence for genetic association between strains causing infection in humans and their pets. Epidemiology & Infection 141, 2526–2535 (2013).

12 Lefebvre, S. L., Reid-Smith, R. J., Waltner-Toews, D. & Weese, J. S. Incidence of acquisition of methicillin-resistant Staphylococcus aureus, Clostridium difficile, and other health-care–associated pathogens by dogs that participate in animal-assisted interventions. Journal of the American Veterinary Medical Association 234, 1404–1417 (2009).

13 López, C. M. et al. Thermotolerant Campylobacters in domestic animals in a defined population in Buenos Aires, Argentina. Preventive Veterinary Medicine 55, 193–200 (2002).

14 Pan, H., Li, W., Sun, E. & Zhang, Y. Characterization and whole genome sequencing of a novel strain of Bergeyella cardium related to infective endocarditis. BMC microbiology 20, 1–9 (2020).

15 Younus, M. et al. The role of exposures to animals and other risk factors in sporadic, non-typhoidal Salmonella infections in Michigan children. Zoonoses and Public Health 57, e170–e176 (2010).

16 Mermin, J. et al. Reptiles, amphibians, and human Salmonella infection: a population-based, case-control study. Clinical Infectious Diseases 38, S253–S261 (2004).

17 Boehm, T. M. & Mueller, R. S. Dermatophytosis in dogs and cats-an update. Tierarztliche Praxis. Ausgabe K, Kleintiere/Heimtiere 47, 257–268 (2019).

18 Elmore, S. A. et al. Toxoplasma gondii: epidemiology, feline clinical aspects, and prevention. Trends in parasitology 26, 190–196 (2010).

19 Borland, S., Gracieux, P., Jones, M., Mallet, F. & Yugueros-Marcos, J. Influenza A virus infection in cats and dogs: a literature review in the light of the “one health” concept. Frontiers in public health 8, 521275 (2020).

20 Weese, J., Finley, R., Reid-Smith, R., Janecko, N. & Rousseau, J. Evaluation of Clostridium difficile in dogs and the household environment. Epidemiology & Infection 138, 1100–1104 (2010).

21 Meyer, E., Gastmeier, P., Kola, A. & Schwab, F. Pet animals and foreign travel are risk factors for colonisation with extended-spectrum β-lactamase-producing Escherichia coli. Infection 40, 685–687 (2012).

22 Kachrimanidou, M., Tzika, E. & Filioussis, G. Clostridioides (Clostridium) difficile in food-producing animals, horses and household pets: a comprehensive review. Microorganisms 7, 667 (2019).

23 Wei, Y. et al. Prevalence, genotype and antimicrobial resistance of Clostridium difficile isolates from healthy pets in Eastern China. BMC Infectious Diseases 19, 1–7 (2019).

24 Loo, V. G., Brassard, P. & Miller, M. A. Household transmission of Clostridium difficile to family members and domestic pets. infection control & hospital epidemiology 37, 1342–1348 (2016).

25 Buranasinsup, S. et al. Prevalence and characterization of antimicrobial-resistant Escherichia coli isolated from veterinary staff, pets, and pet owners in Thailand. Journal of Infection and Public Health 16, 194–202 (2023).

26 Toombs-Ruane, L. J. et al. Carriage of extended-spectrum-beta-lactamase-and AmpC beta-lactamase-producing Escherichia coli strains from humans and pets in the same households. Applied and environmental microbiology 86, e01613–01620 (2020).

27 Xie, L. et al. Analysis of Lung Microbiome in COVID-19 Patients during Time of Hospitalization. Pathogens 12, 944 (2023).

28 Shi, M. et al. Total infectome characterization of respiratory infections in pre-COVID-19 Wuhan, China. PLoS Pathogens 18, e1010259 (2022).

29 Wierenga, J. R. et al. Total infectome investigation of diphtheritic stomatitis in yellow-eyed penguins (Megadyptes antipodes) reveals a novel and abundant megrivirus. Veterinary Microbiology 286, 109895 (2023).

30 Brito, B. P. et al. Expanding the range of the respiratory infectome in Australian feedlot cattle with and without respiratory disease using metatranscriptomics. Microbiome 11, 158 (2023).

31 Cui, X. et al. Virus diversity, wildlife-domestic animal circulation and potential zoonotic viruses of small mammals, pangolins and zoo animals. Nature Communications 14, 2488 (2023).

32 Huang, X. et al. A total infectome approach to understand the etiology of infectious disease in pigs. Microbiome 10, 73 (2022).

33 Wang, J. et al. Emerging sand fly–borne phlebovirus in China. Emerging Infectious Diseases 26, 2435 (2020).

34 Zhong, H. et al. Characterization of respiratory microbial dysbiosis in hospitalized COVID-19 patients. Cell discovery 7, 23 (2021).

35 Jha, A. R. et al. Characterization of gut microbiomes of household pets in the United States using a direct-to-consumer approach. PLoS One 15, e0227289 (2020).

36 Alessandri, G. et al. Metagenomic dissection of the canine gut microbiota: insights into taxonomic, metabolic and nutritional features. Environmental Microbiology 21, 1331–1343 (2019).

37 Moon, C. D., Young, W., Maclean, P. H., Cookson, A. L. & Bermingham, E. N. Metagenomic insights into the roles of Proteobacteria in the gastrointestinal microbiomes of healthy dogs and cats. Microbiologyopen 7, e00677 (2018).

38 Conceição-Neto, N., et al. Viral gut metagenomics of sympatric wild and domestic canids, and monitoring of viruses: Insights from an endangered wolf population. Ecology and Evolution 7, 4135–4146 (2017).

39 Wang, H. et al. Viral Metagenomic Analysis of the Fecal Samples in Domestic Dogs (Canis lupus familiaris). Viruses 15, 685 (2023).

40 Moreno, P. S. et al. Characterisation of the canine faecal virome in healthy dogs and dogs with acute diarrhoea using shotgun metagenomics. PloS one 12, e0178433 (2017).

41 Altan, E. et al. Nasal virome of dogs with respiratory infection signs include novel taupapillomaviruses. Virus genes 55, 191–197 (2019).

42 Zhou, Q., Yu, J., Song, X., Zhang, J. & Zhang, B. The discovery of novel papillomaviruses in cats in Southwest China. Virus Genes 59, 484–488 (2023).

43 Weber, M. et al. Characterization of dog serum virome from Northeastern Brazil. Virology 525, 192–199 (2018).

44 Ng, T. F. F. et al. Feline fecal virome reveals novel and prevalent enteric viruses. Veterinary microbiology 171, 102–111 (2014).

45 Van Brussel, K. et al. The enteric virome of cats with feline panleukopenia differs in abundance and diversity from healthy cats. Transboundary and Emerging Diseases 69, e2952–e2966 (2022).

46 Hoffmann, A. R., Proctor, L., Surette, M. & Suchodolski, J. The microbiome: the trillions of microorganisms that maintain health and cause disease in humans and companion animals. Veterinary pathology 53, 10–21 (2016).

47 Shi, Y. et al. The Gut Viral Metagenome Analysis of Domestic Dogs Captures Snapshot of Viral Diversity and Potential Risk of Coronavirus. Frontiers in Veterinary Science 8, 695088 (2021).

48 Barko, P., McMichael, M., Swanson, K. S. & Williams, D. A. The gastrointestinal microbiome: a review. Journal of veterinary internal medicine 32, 9–25 (2018).

49 McDonald, J. E. et al. Characterising the canine oral microbiome by direct sequencing of reverse-transcribed rRNA molecules. PLoS One 11, e0157046 (2016).

50 Ruparell, A. et al. The canine oral microbiome: variation in bacterial populations across different niches. BMC microbiology 20, 42 (2020).

51 Dewhirst, F. E. et al. The feline oral microbiome: a provisional 16S rRNA gene based taxonomy with full-length reference sequences. Veterinary microbiology 175, 294–303 (2015).

52 Dewhirst, F. E. et al. The canine oral microbiome. PloS one 7, e36067 (2012).

53 Wallis, C. V., Marshall-Jones, Z. V., Deusch, O. & Hughes, K. R. Canine and feline microbiomes. Understanding Host-Microbiome Interactions-An Omics Approach: Omics of Host-Microbiome Association, 279–325 (2017).

54 Lee, S. J. et al. Increasing Fusobacterium infections with Fusobacterium varium, an emerging pathogen. Plos one 17, e0266610 (2022).

55 Smallman, T. R. et al. Pathogenomic analysis and characterization of Pasteurella multocida strains recovered from human infections. Microbiology Spectrum, e03805–03823 (2024).

56 Holst, E., Rollof, J., Larsson, L. & Nielsen, J. P. Characterization and distribution of Pasteurella species recovered from infected humans. Journal of Clinical Microbiology 30, 2984–2987 (1992).

57 Jokelainen, P. et al. Veterinarians and zoonotic pathogens, infections and diseases–questionnaire study and case series, Finland. Infectious Diseases, 1–9 (2024).

58 Holmes, B. et al. Neisseria weaveri sp. nov.(formerly CDC group M-5), from dog bite wounds of humans. International Journal of Systematic and Evolutionary Microbiology 43, 687-693 (1993).

59 Mukherjee, P. K. et al. Mycobiota in gastrointestinal diseases. Nature Reviews Gastroenterology & Hepatology 12, 77–87 (2015). 10.1038/nrgastro.2014.188

60 Poirier, S. et al. Deciphering intra-species bacterial diversity of meat and seafood spoilage microbiota using gyrB amplicon sequencing: A comparative analysis with 16S rDNA V3-V4 amplicon sequencing. PLoS One 13, e0204629 (2018).

61 Scharf, S. et al. Introduction of a bead beating step improves fungal DNA extraction from selected patient specimens. International Journal of Medical Microbiology 310, 151443 (2020).

62 Renshaw, R. W. et al. Characterization of a vesivirus associated with an outbreak of acute hemorrhagic gastroenteritis in domestic dogs. Journal of clinical microbiology 56, 10.1128/jcm.01951-01917 (2018).

63 Charoenkul, K. et al. First detection and genetic characterization of canine Kobuvirus in domestic dogs in Thailand. BMC Veterinary Research 15, 254 (2019). 10.1186/s12917-019-1994-6

64 De Vries, R. D., Duprex, W. P. & De Swart, R. L. Morbillivirus infections: an introduction. Viruses 7, 699–706 (2015).

65 Vlasova, A. N. et al. Novel canine coronavirus isolated from a hospitalized patient with pneumonia in East Malaysia. Clinical Infectious Diseases 74, 446–454 (2022).

66 Villabruna, N., Koopmans, M. P. & de Graaf, M. Animals as reservoir for human norovirus. Viruses 11, 478 (2019).

67 Jadhav, V. J. & Pal, M. Human and domestic animal infections caused by Candida albicans. J. Mycopathol. Res 51, 243–249 (2013).

68 Quast, C. et al. The SILVA ribosomal RNA gene database project: improved data processing and web-based tools. Nucleic acids research 41, D590–D596 (2012).

69 Langmead, B. & Salzberg, S. L. Fast gapped-read alignment with Bowtie 2. Nature methods 9, 357–359 (2012).

70 Fu, L., Niu, B., Zhu, Z., Wu, S. & Li, W. CD-HIT: accelerated for clustering the next-generation sequencing data. Bioinformatics 28, 3150–3152 (2012).

71 Li, D., Liu, C.-M., Luo, R., Sadakane, K. & Lam, T.-W. MEGAHIT: an ultra-fast single-node solution for large and complex metagenomics assembly via succinct de Bruijn graph. Bioinformatics 31, 1674–1676 (2015).

72 Camacho, C. et al. BLAST+: architecture and applications. BMC bioinformatics 10, 1–9 (2009).

73 Buchfink, B., Xie, C. & Huson, D. H. Fast and sensitive protein alignment using DIAMOND. Nature methods 12, 59–60 (2015).

74 Shi, M. et al. The evolutionary history of vertebrate RNA viruses. Nature 556, 197–202 (2018).

75 Parks, D. H. et al. GTDB: an ongoing census of bacterial and archaeal diversity through a phylogenetically consistent, rank normalized and complete genome-based taxonomy. Nucleic acids research 50, D785–D794 (2022).

76 Manni, M., Berkeley, M. R., Seppey, M., Simão, F. A. & Zdobnov, E. M. BUSCO update: novel and streamlined workflows along with broader and deeper phylogenetic coverage for scoring of eukaryotic, prokaryotic, and viral genomes. Molecular biology and evolution 38, 4647–4654 (2021).

77 Kanehisa, M., Furumichi, M., Tanabe, M., Sato, Y. & Morishima, K. KEGG: new perspectives on genomes, pathways, diseases and drugs. Nucleic acids research 45, D353–D361 (2017).

78 Ogier, J.-C., Pagès, S., Galan, M., Barret, M. & Gaudriault, S. rpoB, a promising marker for analyzing the diversity of bacterial communities by amplicon sequencing. BMC microbiology 19, 1–16 (2019).

79 Katoh, K. & Standley, D. M. MAFFT multiple sequence alignment software version 7: improvements in performance and usability. Molecular biology and evolution 30, 772–780 (2013).

80 Minh, B. Q. et al. IQ-TREE 2: new models and efficient methods for phylogenetic inference in the genomic era. Molecular biology and evolution 37, 1530–1534 (2020).

81 Van der Maaten, L. & Hinton, G. Visualizing data using t-SNE. Journal of machine learning research 9 (2008).

82 McArdle, B. H. & Anderson, M. J. Fitting multivariate models to community data: a comment on distance-based redundancy analysis. Ecology 82, 290–297 (2001).

83 Love, M. I., Huber, W. & Anders, S. Moderated estimation of fold change and dispersion for RNA-seq data with DESeq2. Genome biology 15, 1–21 (2014).

84 Segata, N. et al. Metagenomic biomarker discovery and explanation. Genome biology 12, 1–18 (2011).

85 Nearing, J. T. et al. Microbiome differential abundance methods produce different results across 38 datasets. Nature communications 13, 342 (2022).

