## Supplementary material for "Meta-transcriptomic analysis of companion animal infectomes reveals their diversity and potential roles in animal and human disease": Suppmental Figure S1-S7

**Supplementary Figures**

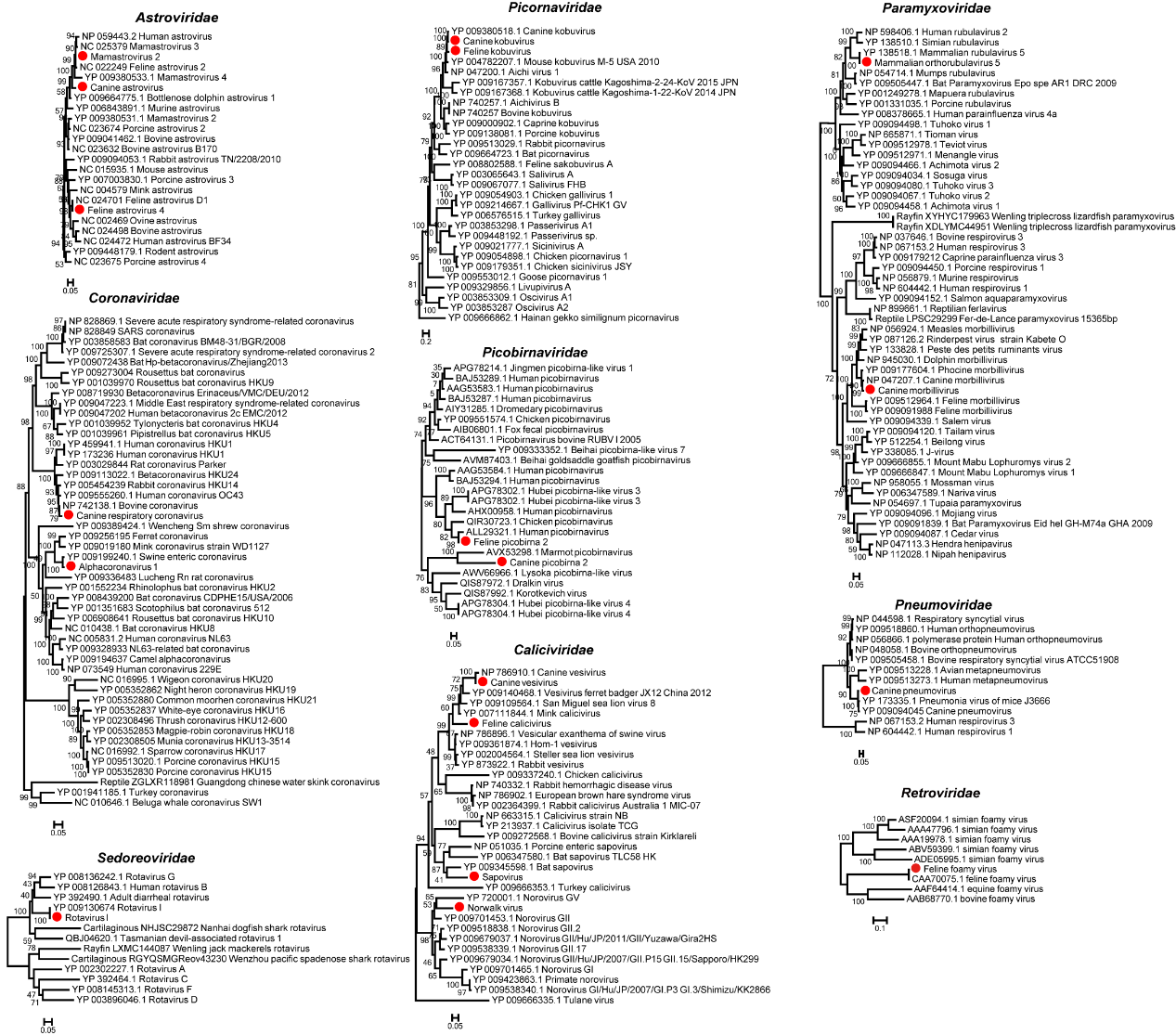

**Figure S1. Maximum likelihood phylogenetic trees of RNA viruses.** The phylogenies were inferred based on multiple sequence alignments of the RNA-dependent RNA polymerase (RdRp) protein, with the exception of the *Retroviridae* for which the reverse transcriptase (RT) protein was used The trees are midpoint rooted for clarity, with branch lengths reflecting the number of substitutions per site.

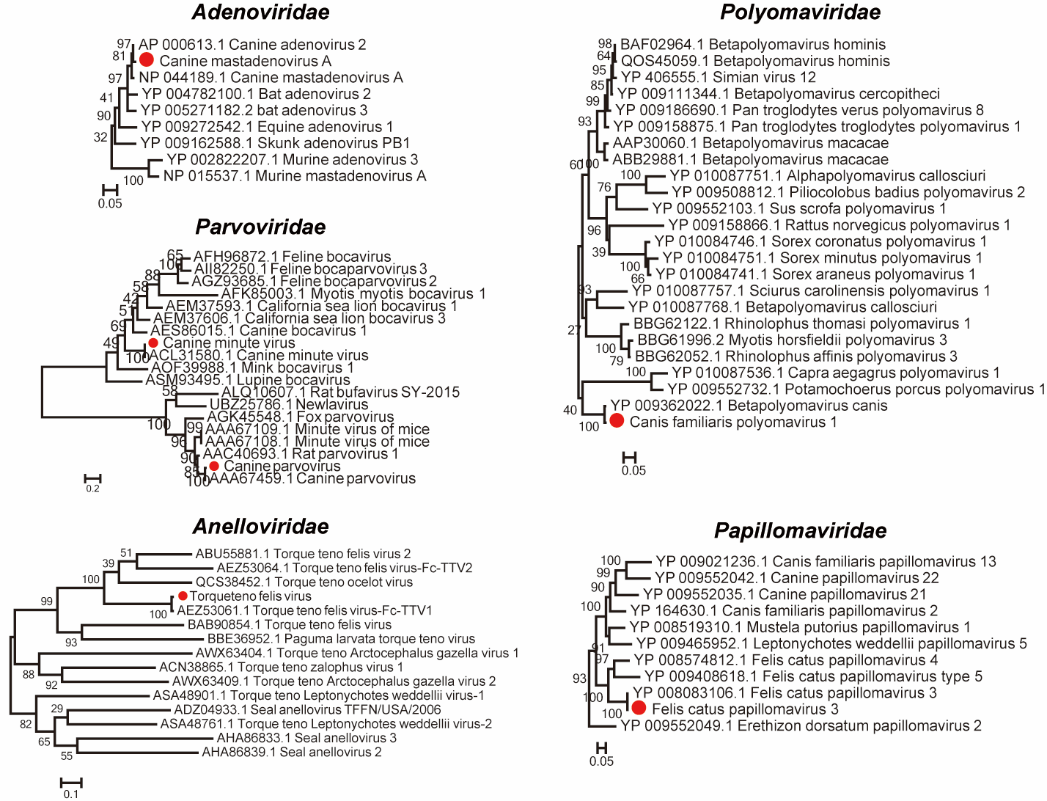

**Figure S2. Maximum likelihood phylogenetic trees of DNA viruses.** The phylogenies were inferred using DNA polymerase protein (*Adenoviridae*), NS1 protein (*Parvoviridae*), ORF1 protein (*Anelloviridae*), E1 protein (*Papillomaviridae*), and LT-Ag protein (*Polyomaviridae*). The trees are midpoint rooted for clarity, with branch lengths reflecting the number of substitutions per site.

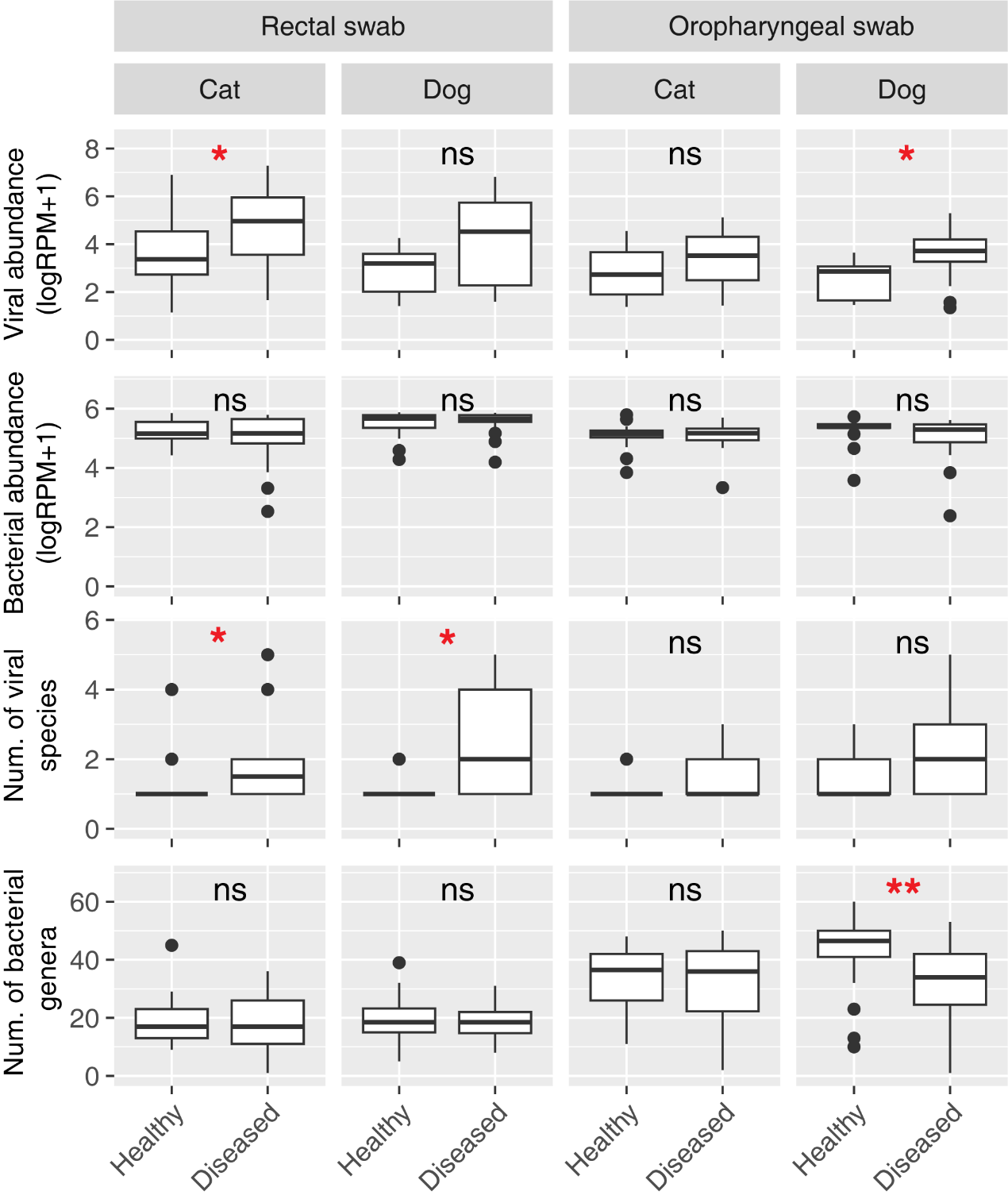

**Figure S3. Comparisons of viral and bacterial abundance and diversity in healthy and diseased animals.** Statistical significance was determined using two-sided Wilcoxon rank-sum tests. ns: not significant (p > 0.05), *: 0.01 < p ≤ 0.05, **: 0.001 < p ≤ 0.01.

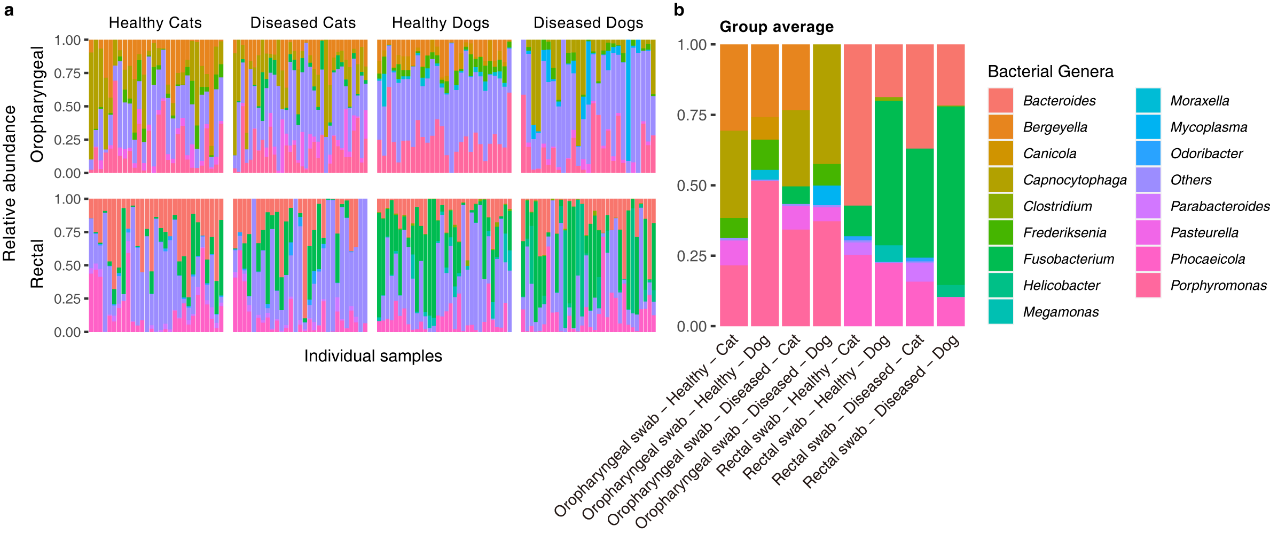

**Figure S4. Bacterial community composition in healthy and diseased cats and dogs.** (a) Bacterial genera composition in each sample. (b) Bar plot showing the average relative abundances of major bacterial genera within each group. The y-axis represents the relative abundance, and the x-axis labels denote the different groups. Colors are used to distinguish different bacterial genera (including top 5 most abundant genera in of group).

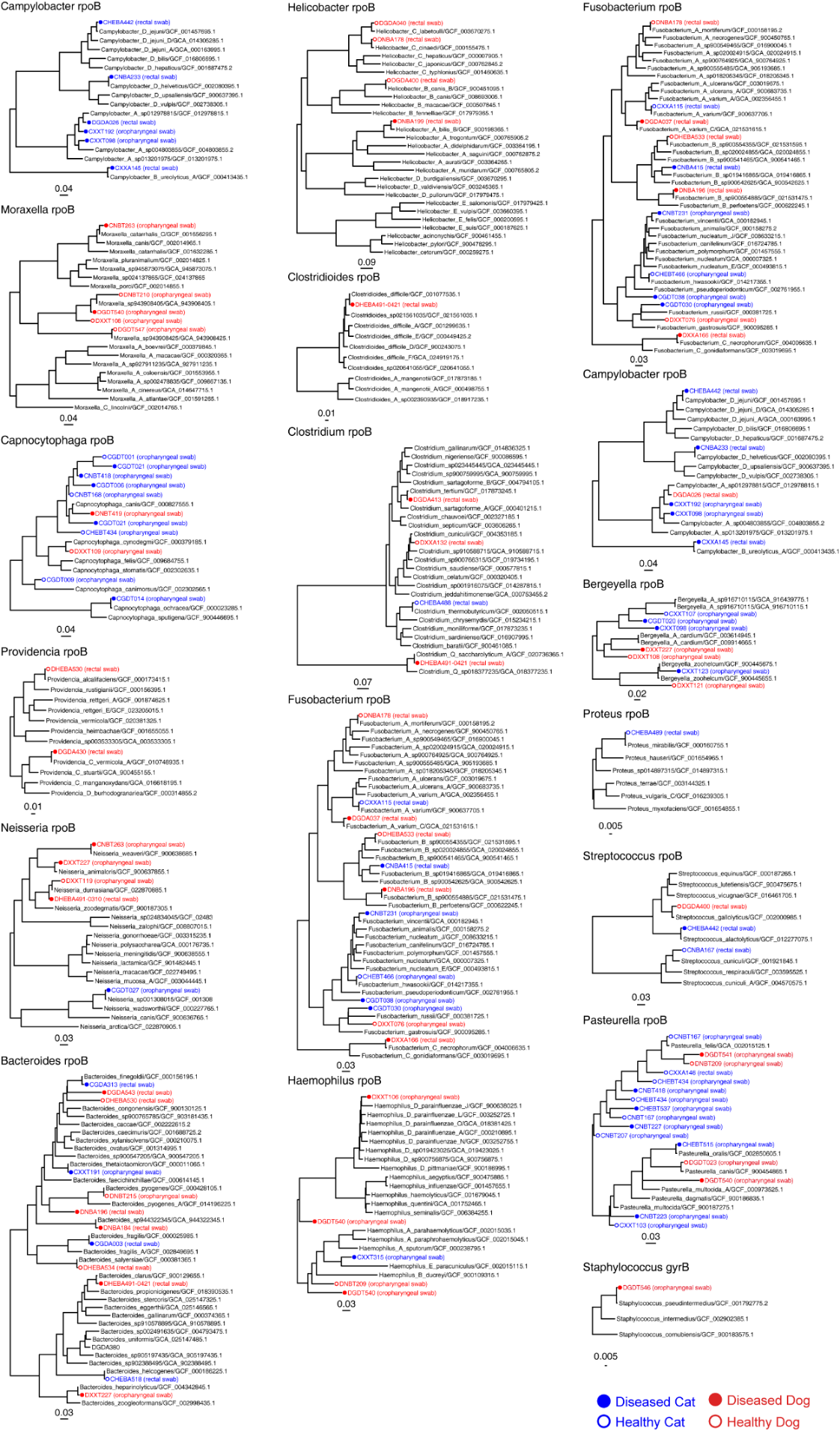

**Figure S5. Maximum likelihood phylogenetic trees of potential zoonotic bacterial species.** The phylogenies were inferred using nucleotide sequences of rpoB gene. The trees are midpoint rooted for clarity, with branch lengths reflecting the number of substitutions per site.

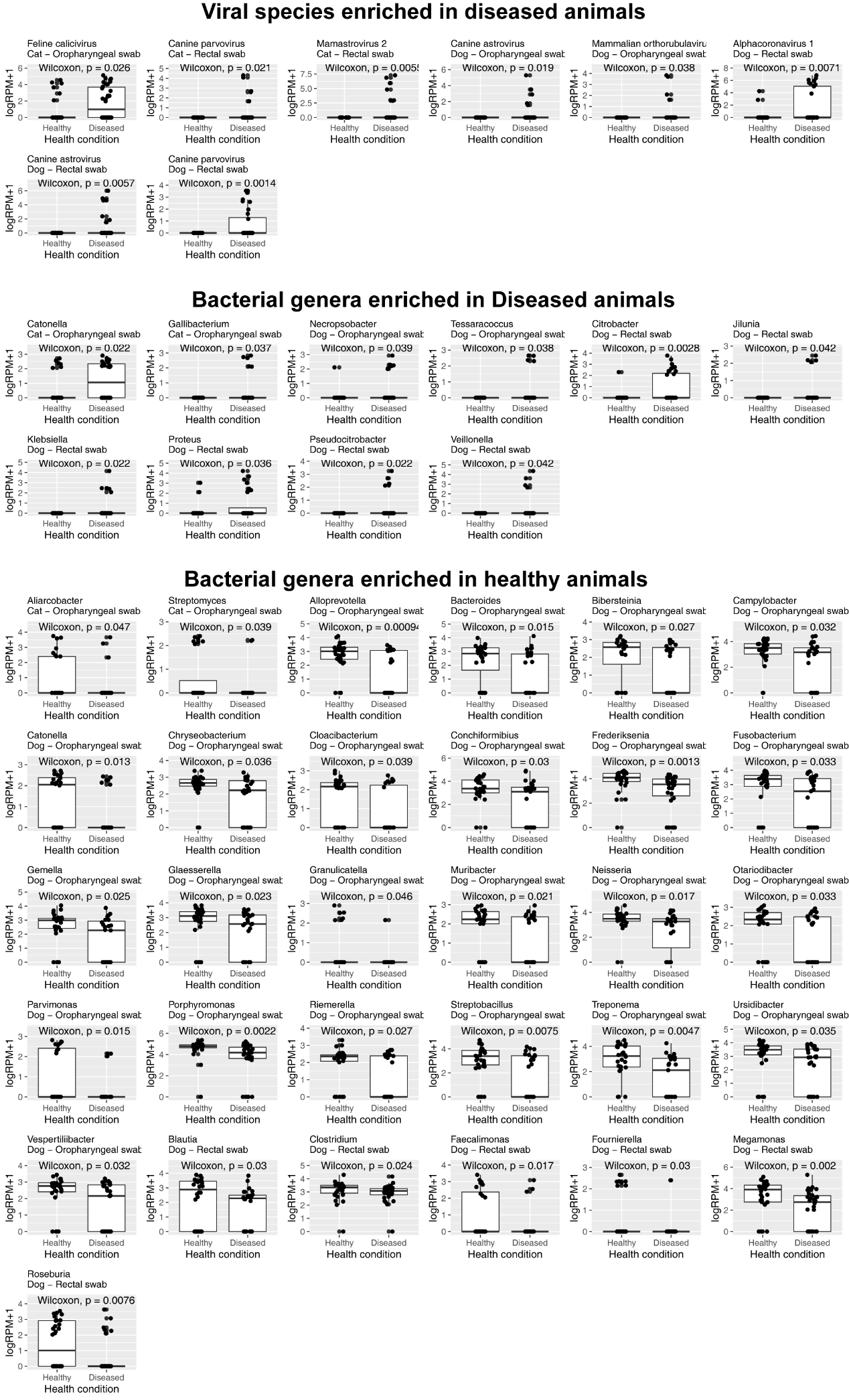

**Figure S6. Viral species and bacterial genera enriched in diseased and healthy animals**. The top panel displays box plots comparing the relative abundances of different viral species between healthy and diseased animals. The middle and bottom panels show similar box plots for bacterial genera that are enriched in diseased and healthy animals, respectively. Each subplot is labeled with the taxonomic name and includes a p-value (Wilcoxon tests) indicating the statistical significance of the difference between the two groups.

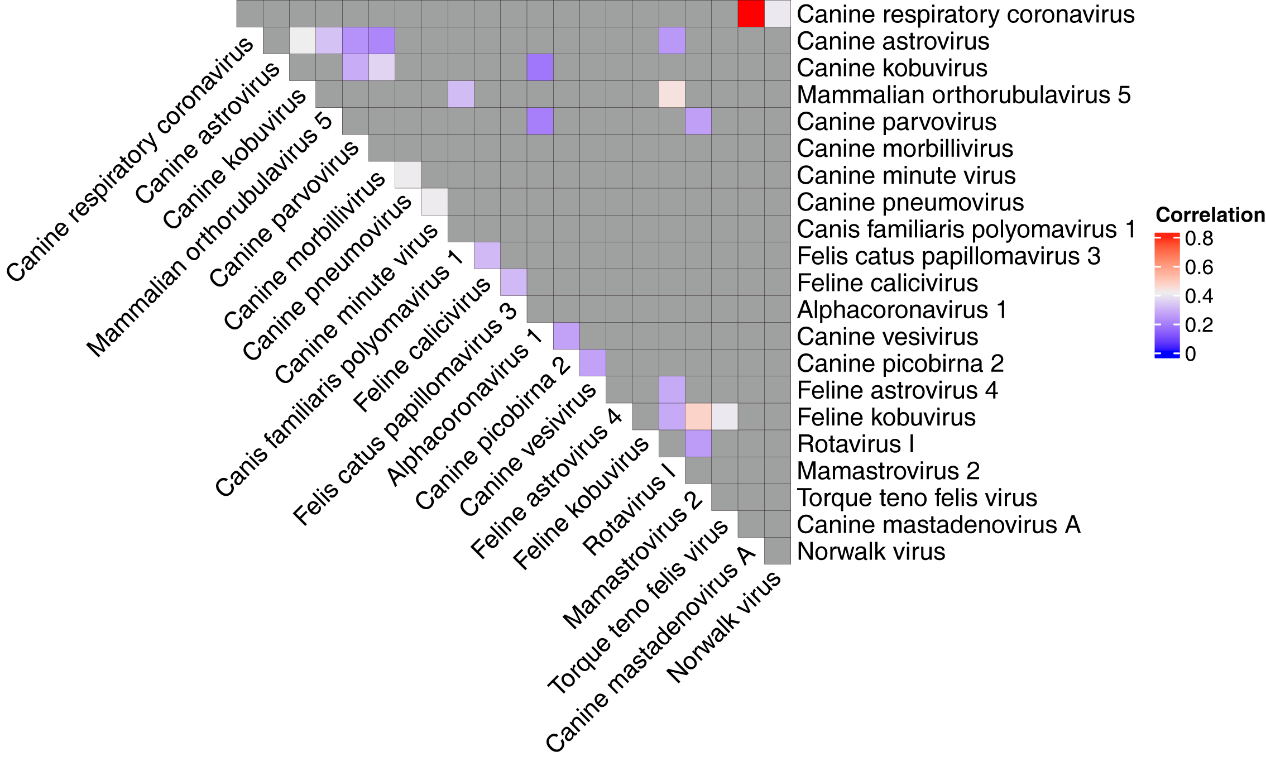

**Figure S7. Heatmap illustrating virus co-infections in companion animals.** Colors represent the Spearman correlation between the abundance level of two viruses. All the significant correlations (p < 0.05) tested here are positive, with the red color indicating a stronger positive correlation. The gray color indicates non-significant correlations.
